# Time preferences are reliable across time-horizons and verbal vs. experiential tasks

**DOI:** 10.1101/351312

**Authors:** Evgeniya Lukinova, Yuyue Wang, Steven F. Lehrer, Jeffrey C. Erlich

## Abstract

Individual differences in delay-discounting correlate with important real world outcomes, e.g. education, income, drug use, & criminality. As such, delay-discounting has been extensively studied by economists, psychologists and neuroscientists to reveal its behavioral and biological mechanisms in both human and non-human animal models. However, two major methodological differences hinder comparing results across species. Human studies present long time-horizon options verbally, whereas animal studies employ experiential cues and short delays. To bridge these divides, we developed a novel language-free experiential task inspired by animal decision-making studies. We find that subjects’ time-preferences are reliable across both verbal/experiential differences and also second/day differences. When we examined whether discount factors shifted or scaled across the tasks, we found a surprisingly strong effect of temporal context. Taken together, this indicates that subjects have a stable, but context-dependent, time-preference that can be reliably assessed using different methods; thereby, providing a foundation to bridge studies of time-preferences across species.

## Introduction

Intertemporal choices involve a trade-off between a larger outcome received later and a smaller outcome received sooner. Many individual decisions have this temporal structure, such as whether to purchase a cheaper refrigerator, but forgo the ongoing energy savings. Since research has found that intertemporal preferences are predictive of a wide variety of important life outcomes, ranging from SAT scores, graduating from college, and income to anti-social behaviors, e.g. gambling or drug abuse [1,13,24,28,45], they are frequently studied in both humans and animals across multiple disciplines, including marketing, economics, psychology, and neuroscience.

A potential obstacle to understanding the biological basis of intertemporal decision-making is that human studies differ from non-human animal studies in two important ways: long versus short time-horizons and choices that are made based on verbal versus non-verbal (i.e. “experiential”) stimuli. In animal studies, the subjects experience the delay between their choice and the reward (sometimes cued with a ramping sound or a diminishing visual stimulus) before they can proceed to the next trial [8,11,73]. Generally, there is nothing for the subject to do during this waiting period. In human studies, subjects usually make a series of choices (either via computer or a survey) between smaller sooner offers and larger offers delayed by months or years [2,49]. (We are aware of only a handful of studies that have used delays of minutes [48] or seconds [25,31,40,60,72]). During the delay (e.g. if the payout is in 6 months) the human subjects go about their lives, likely forgetting about the delayed payment, just as individuals do not actively think about their retirement savings account each moment until their retirement.

Animal studies of delay-discounting take several forms [11,18,62,79], but all require experiential learning that some non-verbal cue is associated with waiting. Subjects experience the cues, delays and rewards, and slowly build an internal map from the cues to the delays and magnitudes. Subjects may only have implicit knowledge of the map, which likely engage distinct neural substrates to the explicit processes engaged by humans when considering a verbal offer [59,61].

Whether animal studies can inform human studies depends on answers to the following questions. Do decisions that involve actively waiting for seconds invoke the same cognitive and neural processes as decisions requiring passively waiting for months? Do decisions made based on experience and perceptual decisions invoke the same cognitive and neural processes as decisions that are made based on explicitly written information?

The animal neuroscience literature on delay-discounting mostly accepts as a given that the behavior of animals will give insight into the biological basis for human impulsivity [23,34,65,70] and rarely [8,66] addresses the methodological gaps considered here. This view is not unfounded. Neural recordings from animals [11] and brain imaging studies in humans [37,49] both find that the prefrontal cortex and basal ganglia are involved in delay-discounting decisions, suggesting common neural mechanisms. Animal models of attention-deficit hyperactive disorder (ADHD) have reasonable construct validity: drugs that shift animal behavior in delay-discounting tasks can also improve the symptoms of ADHD in humans [23,56]. Thus, most neuroscientists would likely predict that our experiments would find high within-subject reliability across both time-horizons and verbal/experiential dimensions.

Reading the literature from economics, a different picture emerges. Traditional economic models dating back to Samuelson [68] posit that agents make consistent intertemporal decisions, thereby implying a constant discount rate regardless of context. In contrast, growing evidence from behavioral economics provides support for the view that discounting over a given time delay changes with the time-horizon [3,7]. Yet, there remains debate in the empirical economics literature about how well discounting measures elicited in human studies truly reflect the rates of time-preference used in real-world decisions since they have been found to vary by the type of task (hypothetical, potentially real, and real), stakes being compared, age of participants and across different domains [15]. Thus, most economists surveying the empirical evidence would be surprised if a design that varied both type of tasks and horizons would generate results with high within-subject reliability.

Here, we have addressed these questions by measuring the discount factors of human subjects in three ways. First, we used a novel language-free task involving experiential learning with short delays [20,42,43,46]. Then, we measured discount factors more traditionally, with verbal offers over both short and long delays. This design allowed us to test whether, for each subject, a single process is used for intertemporal choice regardless of time-horizon or verbal vs. experiential stimuli, or whether the choices in different tasks could be better explained by distinct underlying mechanisms.

## Results

In our main experiment, 63 undergraduate students from NYU Shanghai participated in 5 experimental sessions. In each session, subjects completed a series of intertemporal choices. Across sessions, 160 trials were conducted involving each of the following 3 tasks, i) non-verbal short delay (NV, 3 - 64 seconds), ii) verbal short delay (SV, 3 - 64 seconds), and iii) verbal long delay (LV, 3 - 64 days). In each trial, irrespective of the task, subjects made a decision between the sooner (blue circle) and the later (yellow circle) options. In the non-verbal task (Fig. 1A) the parameters of the later option were mapped to an amplitude modulated pure tone. The reward magnitude was mapped to frequency of the tone (larger reward ∝ higher frequency). The delay was mapped to amplitude modulation rate (longer delay ∝ slower modulation). Across trials, the delay and the magnitude of the sooner option were fixed (4 coins, immediately). For the short delay tasks, when subjects chose the later option, a clock appeared on the screen, and only when the clock image disappeared, could they collect their reward by clicking in the reward port. The rewards were accumulated for the duration of the task and used for subject’s payment. In the verbal tasks, the verbal description of the offers appeared within the blue & yellow circles in place of the amplitude modulated sound (Fig. 1B). In the verbal long delay task, after each choice, subjects were given feedback confirming their choice and then proceeded to the next trial. At the end of the session, a single long-verbal trial was selected randomly to determine the payment. If the selected trial corresponded to a subject having chosen the later option, she received her reward via an electronic transfer after the delay.

**Figure 1.**
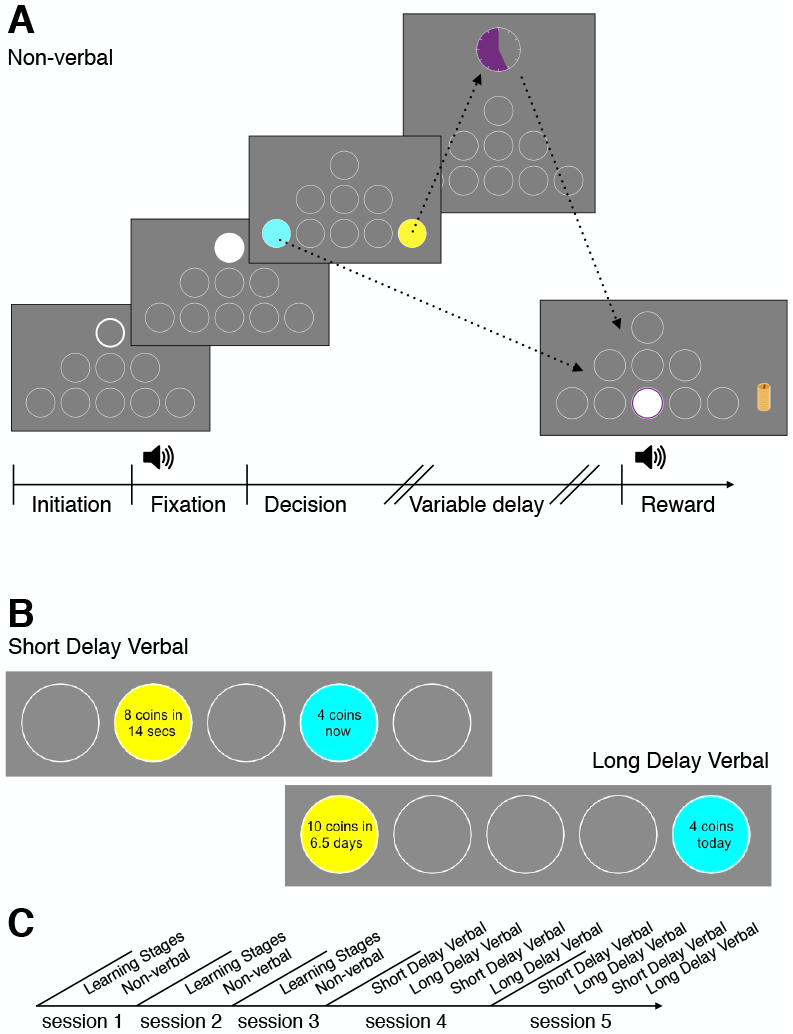
A: A novel language-free intertemporal choice task. This is an example sequence of screens that subjects viewed in one trial of the non-verbal task. First, the subject initiates the trial by pressing on the white-bordered circle. During fixation the subject must keep the cursor inside the white circle. The subject hears an amplitude modulated pure tone (the tone frequency is mapped to reward magnitude and the modulation rate is mapped to the delay of the later option). The subject next makes a decision between the sooner (blue circle) and later (yellow circle) options. If the later option is chosen, the subject waits until the delay time finishes - which is indicated by the colored portion of the clock image. Finally, the subject clicks in the middle bottom circle (”reward port”) to retrieve their reward. The reward is presented as a stack of coins of a specific size and a coin drop sound accompanies the presentation. B: Stimuli examples in the verbal experiment during decision stage (the bottom row of circles is cropped). C: Timeline of experimental sessions.

### Subjects’ time-preferences are reliable across both verbal/experiential and second/day differences

Subjects’ impulsivity was estimated by fitting their choices with a hierarchical Bayesian model of hyperbolic discounting with decision noise (*Materials and Methods*). The model (*M*_6*p*,4*s*_) had 6 population level parameters (discount factor, *k*, and decision noise, *τ*, for each of the three tasks) and 4 parameters per subject: *k*_*NV*_, *k*_*SV*_,*k*_*LV*_ and τ. The subject level effects are drawn from a normal distribution with mean zero. Subjects’ choices were well-fit by the model (Fig. 2 & Fig. S1). Since we did not *ex ante* have a strong hypothesis about how the subjects’ impulsivity measures in one task would translate across tasks, we first examined ranks of impulsivity and found significant correlations across experimental tasks (Table 1). In other words, the most impulsive subject in one task is likely to be the most impulsive subject in another task. This result is robust to different functional forms of discounting and estimation methods (Fig. S2 & Table S3). For example, if we ranked the subjects by the fraction of trials they chose the later option in each task, we obtain a similar result (Spearman *r*: SV vs. NV *r* = 0.71; SV vs. LV *r* = 0.49; NV vs. LV *r* = 0.30, all *p* < 0.05) (See *SI Results* for additional confirmations). The correlations of discount factors across tasks extended to Pearson correlation of *log*(*k*) (Fig. 3 & Table 1). We found that *k*, for all tasks, had a log-normal distribution across our subjects (as in [69]) and shown in Fig. 3C), hence we present our results in *log*(*k*).

**Table 1.** Correlations of subjects’ discount factors

|  | Spearman<br>Rank Correlation | Pearson<br>Correlation |
| --- | --- | --- |
| SV vs. NV | 0.76 | 0.79 |
| SV vs. LV | 0.54 | 0.61 |
| NV vs. LV | 0.36 | 0.40 |
| all $p < 0.01$ | | |

Consistent with existing research, we find that time-preferences are stable in the same task within subjects between the first half of the block and the second half of the block within sessions and also across experimental sessions that take place every two weeks (*SI Results*) [4,50]. In our verbal experimental sessions the short and long tasks were alternated and the order was counter-balanced across subjects. We did not find any order effects (*Materials and Methods*) in both main (bootstrapped mean test, SV-LV-SV-LV vs. LV-SV-LV-SV order for SV and LV *log*(*k*), respectively, all p ¿ 0.4) and control experiments (*SI Results*).

**Figure 2.**
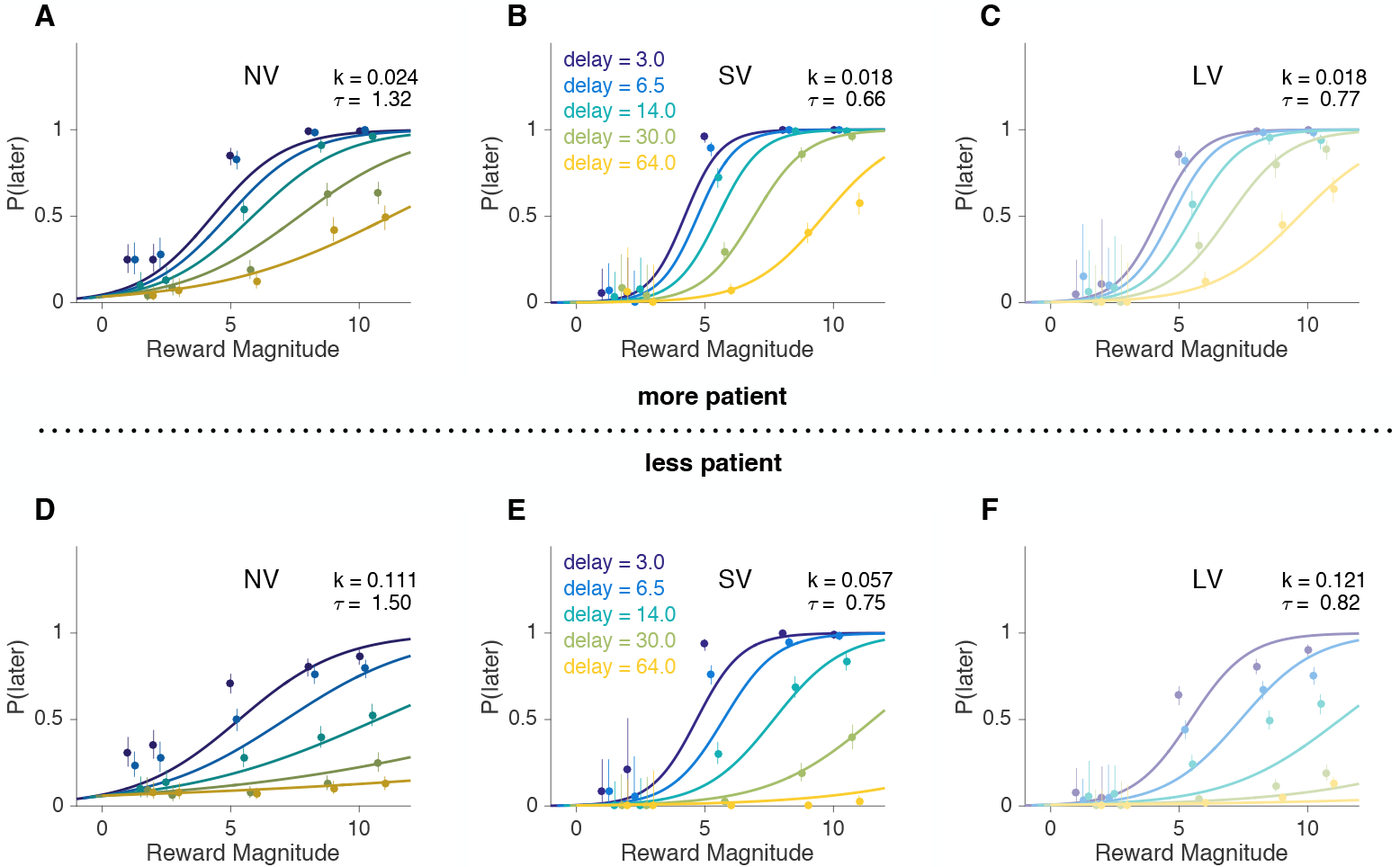
A 50% median split (± 1 standard deviation) of the softmax-hyperbolic fits for more patient (A-C) and less patient (D-F) subjects. The values of *k* and *τ* are the means within each group (decision noise *τ* decreases significantly from non-verbal to verbal tasks, bootstrapped mean test, *p* < 10^−4^). Average psychometric curves obtained from the model fits (lines) versus actual data (circles with error bars) for NV, SV and LV tasks for each delay value, where the x-axis is the reward magnitude and the y-axis is the probability (or proportion for actual choices) of later choice. Error bars are binomial 95% confidence intervals. We excluded the error in the model for visualization.

In our experimental design, the SV task has shared features with both the NV and LV task. First, the SV shares time-horizon with the NV task. Second, the SV and LV are both verbal and were undertaken at the same time. The NV and LV tasks differ in both time-horizon and verbal/non-verbal. The only potential feature that is shared between all tasks is delay-discounting. To test whether the correlation between NV and LV might be accounted for by their shared correlation with the SV task, we performed linear regressions of the discount factors in each task as a function of the other tasks (e.g. *log*(*k*_*NV*_) = *β*_*SV*_*log*(*k*_*SV*_) + *β*_*LV*_*log*(*k*_*LV*_ + *β*_0_ + *ϵ*)). For *NV* the two predictors explained 63% of the variance (*F*(60, 2) = 50.63, *p* < 10^−9^). It was found that *log*(*k*_*SV*_) significantly predicted *log*(*k*_*NV*_) (*β*_*SV*_ = 1.28 ± 0.15, *p* < 10^−9^) but *log*(*k*_*LV*_) did not (*β*_*LV*_ = −0.12 ± 0.09, *p* = 0.181). For *LV* we were able to predict 40% of the variance (*F*(60, 2) = 19.64, *p* < 10^−6^) and found that *log*(*k*_*SV*_) significantly predicted *log*(*k*_*LV*_) (*β*_*SV*_ = 1.26 ± 0.26, *p* < 10^−5^) but *log*(*k*_*NV*_) did not (*β*_*NV*_ = −0.24 ± 0.18, *p* = 0.181). For *SV* the two predictors explained 72% of the variance (*F*(60, 2) = 78.93, *p* < 10^−9^). Coefficients for both predictors were significant (*β*_*NV*_ = 0.435 ± 0.050, *p* < 10^−9^; *β*_*LV*_ = −0.223 ± 0.046, *p* < 10^−5^); where *β* = *mean* ± *std.error*. We further verified these results by generating 1-predictor reduced models based on the stronger of the 2-predictors for each task and comparing the nested models using Akaike Information Criteria (AIC) and likelihood ratio tests (LR test) (Table 2).

**Figure 3.**
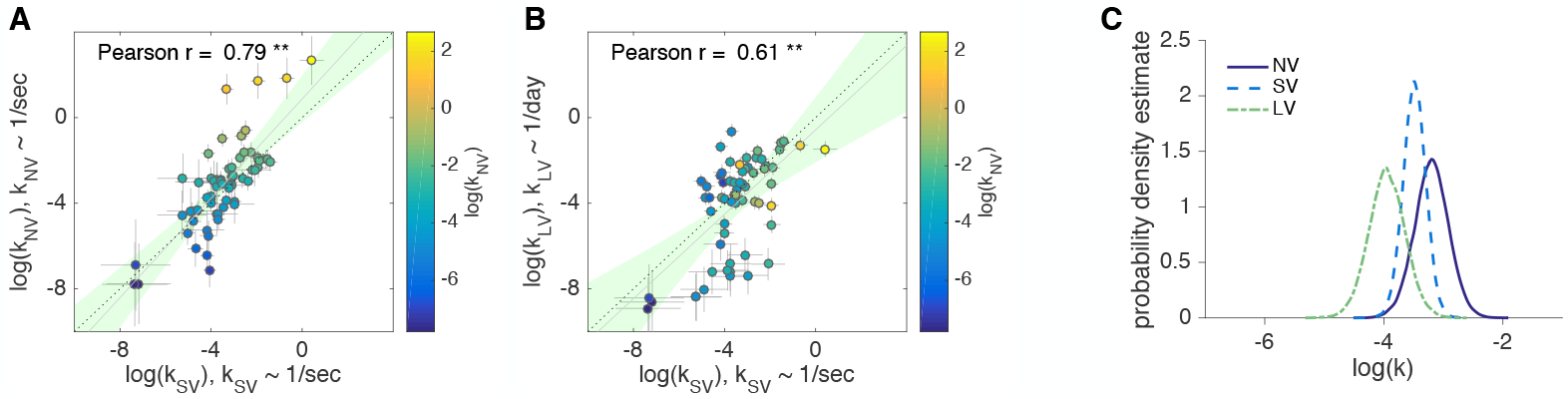
A, B: Each circle is one subject (N=63). The logs of delay-discounting coefficients in SV task (x-axis) plotted against the logs of delay-discounting coefficients in NV (A) and LV (B) tasks (y-axis). The color of the circles and the colorbar identify the ranks in NV task. Pearson’s r is reported on the figure. The error bars are the SD of the estimated coefficients (posterior means). Solid line is the linear fit. Shaded area is the 95% CI of the linear fit. Dashed line is unity. C: Distribution of posterior parameter estimates (*log*(*k*)) from the model fit for the three tasks in control experiment 1. Probability density estimates were obtained (*Materials and Methods*)) for posteriors of *log*(*k*) for experimental tasks (*k*_*NV*_ ~ 1/*sec*, *k*_*SV*_ ~ 1/*sec*, *k*_*LV*_ ~ 1/*day*). Comparisons between tasks are reported in Table 4. Note, the units for *k*_*SV*_ & *k*_*NV*_ (1/*sec*) would need to be scaled by (86400*secs*/*day*) to be directly compared to *k*_*LV*_.

**Table 2.**
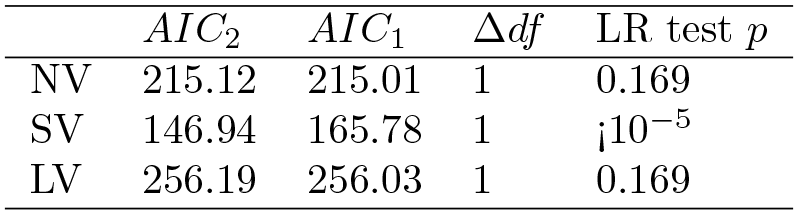
Comparison of 1 vs. 2 predictor linear models of *log*(*k*)

In order to test whether the verbal/non-verbal gap or the time-horizons gap accounted for more variation in discounting we used a linear mixed-effects model where we estimated *log*(*k*) as a function of the two gaps (as fixed effects) with subject as a random effect (using the **lme4** R package [5,6]). We created two predictors: *days* was false in NV and SV tasks for offers in seconds and was true in the LV task for offers in days; *verbal* was true for the SV and LV tasks and false for the NV task. We found that time-horizon (*β*_*days*_ = −0.524 ± 0.235, *p* = 0.026) but not verbal/non-verbal (*β*_*verbal*_ = −0.317 ± 0.235, *p* = 0.178) contributed significantly to the variance in *log*(*k*). This result was further supported by comparing the 2-factor model with reduced 1-factor models (i.e. that only contained either time or verbal fixed effects). Dropping the *days* factor significantly decreased the likelihood, but dropping the *verbal* factor did not (Table 3).

**Table 3.** Relative contributions of two gaps to variance in *log*(*k*)

| Dropped Factor | $\Delta df$ | AIC | LR test $p$ |
| --- | --- | --- | --- |
| *none* |  | 743.06 |  |
| verbal | 1 | 742.88 | 0.177 |
| days | 1 | 745.99 | 0.026 |

We found that subject’s time-preferences were highly correlated across tasks. However, correlation is invariant to shifts or scales across tasks. Our hierarchical model allows us to directly estimate the posterior distributions of *log*(*k*) (Fig. 3C) and report posterior means and credible intervals (NV posterior mean = -3.2, 95% credible interval [-3.77, -2.64], SV mean = -3.49, 95% credible interval [-3.86, -3.11], LV mean = -3.95, 95% credible interval [-4.55, -3.34]). Similarly, we can use the posterior probability to test if *log*(*k*) shifted and/or scaled between tasks (*Materials and Methods*). We find that subjects in both SV and NV are more impatient than LV, but not different from each other (i.e. significant shifts between SV and LV, NV and LV, Table 4). There is significant scaling between SV and the other two tasks (Table 4). This is likely driven by subgroups that were exceptionally patient in the LV task (Fig. 3B) or impulsive in the NV task (Fig. 3A).

**Table 4.** Shift and scale of *log*(*k*) between tasks

| Comparison | $\log_2(\text{Ev. Ratio})$ |
| --- | --- |
| $\mu \log(k_{SV}) > \mu \log(k_{LV})$ | 4.97 * |
| $\mu \log(k_{NV}) > \mu \log(k_{LV})$ | 6.43 * |
| $\mu \log(k_{NV}) > \mu \log(k_{SV})$ | 3.76 |
| $\sigma \log(k_{LV}) > \sigma \log(k_{SV})$ | 7.19 * |
| $\sigma \log(k_{NV}) > \sigma \log(k_{LV})$ | 1.36 |
| $\sigma \log(k_{NV}) > \sigma \log(k_{SV})$ | 12.29 * |

\* denotes $p < 0.05$ one-sided test

### Controlling for visuo-motor confounds

In the main experiment, we held the following features constant across three tasks: the visual display and the use of a mouse to perform the task. However, after observing the strong correlations between the tasks (Fig. 3) we were concerned that the effects could have been driven by the superficial (i.e. visuo-motor) aspects of the tasks. In other words, the visual and response features of the SV and LV tasks may have reminded subjects of the NV task context and nudged them to use a similar strategy across tasks. While this may be interesting in its own right, it would limit the generality of our results. To address this, we ran a control experiment (n=25 subjects) where the NV task was identical to the original NV task, but the SV and LV tasks were run in a more traditional way, with a text display and keypress response (control experiment 1, *SI Method* & Fig. S6). We replicated the main findings of our original experiment for ranks of *log*(*k*) (Table S5) and correlation between *log*(*k*) in SV and LV tasks (Fig. 4B). The Pearson correlation between NV and SV tasks (Fig. 4A) was lower than expected given the 95% confidence intervals of the resampled correlations of the main experiment and assuming 25 subjects (*SI Results*). This suggests that some of the correlation between SV and NV tasks in the main experiment may be driven by visuo-motor similarity in experimental designs. We did not find shifts or scaling between the posterior distributions of *log*(*k*) across tasks in this control experiment (Fig. 4C, NV posterior mean = -3.98, 95% credible interval [-5.44, -2.67], SV mean = -3.8, 95% credible interval [-4.94, -2.75], LV mean = -3.76, 95% credible interval [-4.79, -2.76]).

**Figure 4.**
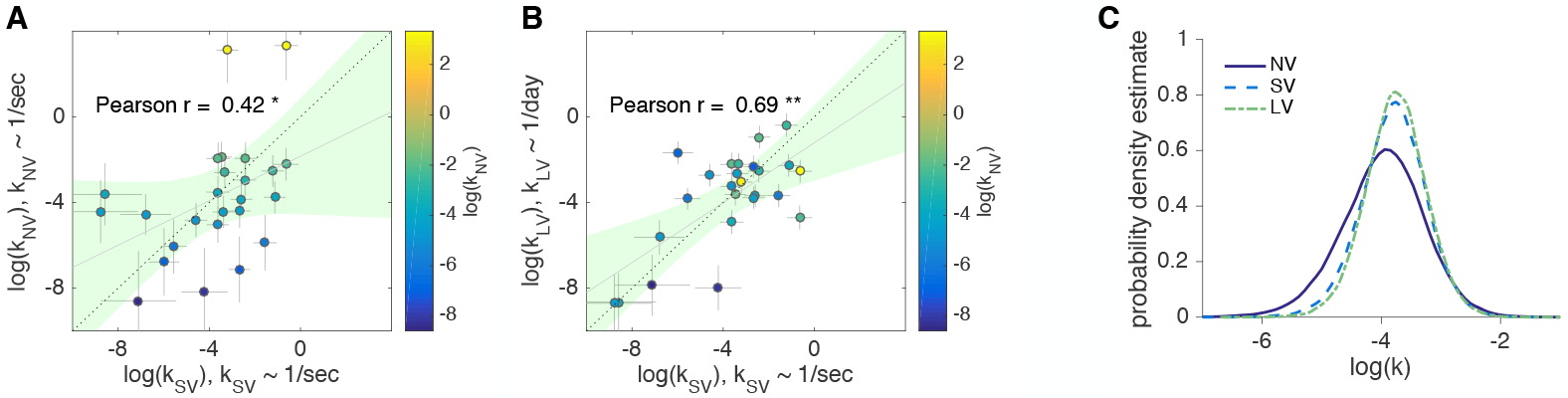
A,B: Control experiment 1 (n=25). The logs of delay-discounting coefficients in SV task (x-axis) plotted against the logs of delay-discounting coefficients in NV (A) and LV (B) tasks (y-axis). The color of the circles and the colorbar identify the ranks in NV task. Each circle is one subject. Pearson’s r is reported on the figure. The error bars are the SD of the estimated coefficients. Solid line is the linear fit. Shaded area is the 95% CI of the linear fit. Dashed line is unity. C: Distribution of posterior parameter estimates (*log*(*k*)) from the model fit for the three tasks in control experiment 1. Probability density estimates were obtained for posteriors of *log*(*k*) for experimental tasks (*k*_*NV*_ ~ 1/*sec*, *k*_*SV*_ ~ 1/*sec*, *k*_*LV*_ ~ l/*day*).

### Strong effect of temporal context

We described above that the discount factors in the LV task, *k*_*LV*_, were almost equivalent (ignoring unexplained variance) to those in the SV task *k*_*SV*_ (Fig. 3B). However, the units of *k*_*LV*_ are in 1/*day* and the units of *k*_*SV*_ are in 1/*seconds*. This finding implies that for a specific reward value, if a subject would decrease their subjective utility of that reward by 50% for an increase from 5 to 10 seconds in the SV task, they would also decrease their subjective utility of that reward by 50% for an increase from 5 to 10 *days* in the LV task. This seems implausible, particularly from a neoclassical economics perspective. However, reward units also change when moving from SV to LV task. In our sessions, the exchange rate in SV was 0.05 CNY per coin (since all coins are accumulated and subjects are paid the total profit), whereas in LV, subjects were paid on the basis of a single trial chosen at random using an exchange rate of 4 CNY for each coin. These exchange rates were set to, on average, equalize the possible total profit between short and long delays tasks. However, even accounting for both the magnitude effect [29,30] and unit conversion (calculations presented in *SI Results*) the discount rates are still scaled by 4 orders of magnitude from the short to the long time-horizon tasks [53].

One interpretation of this result is that subjects are simply ignoring the units and only focusing on the number. This would be consistent with an emerging body of evidence that numerical value, rather than conversion rate or units matter to human subjects [17,26]. A second possible interpretation is that subjects normalize the subjective delay of the offers based on context, just as they normalize subjective value based on current context and recent history [39,41,76,78]. A third possibility is that in the short delay tasks (NV and SV) subjects experience the wait for the reward on each trial as quite costly, in comparison to the delayed gratification experienced in the LV task. This “cost of waiting” may share some intersubject variability with delay-discounting but may effectively scale the discount factor in tasks with this feature [55].

In an attempt to disentangle these possibilities, we ran a control experiment (n=16 subjects) using two verbal discounting tasks (control experiment 2, *SI Method*). In one task, the offers were in days (DV). In the other, the offers were in weeks (WV). This way, we could directly test whether subjects would discount the same for 1 day as 1 week (i.e. ignore units) or 7 days as 1 week (i.e. convert units). We found strong evidence for the latter (Fig. 5A). Subjects did not ignore the units: their choices were consistent with rational agents that converted all offers into a common time unit. There is almost perfect correlation (Pearson *r* = 0.97, *p* < 0.01) across estimated *log*(*k*) (*k* ~ 1/*day*) within subjects between verbal task with delays in days and delays in weeks (see control experiment 2 results in *SI Results*).

**Figure 5.**
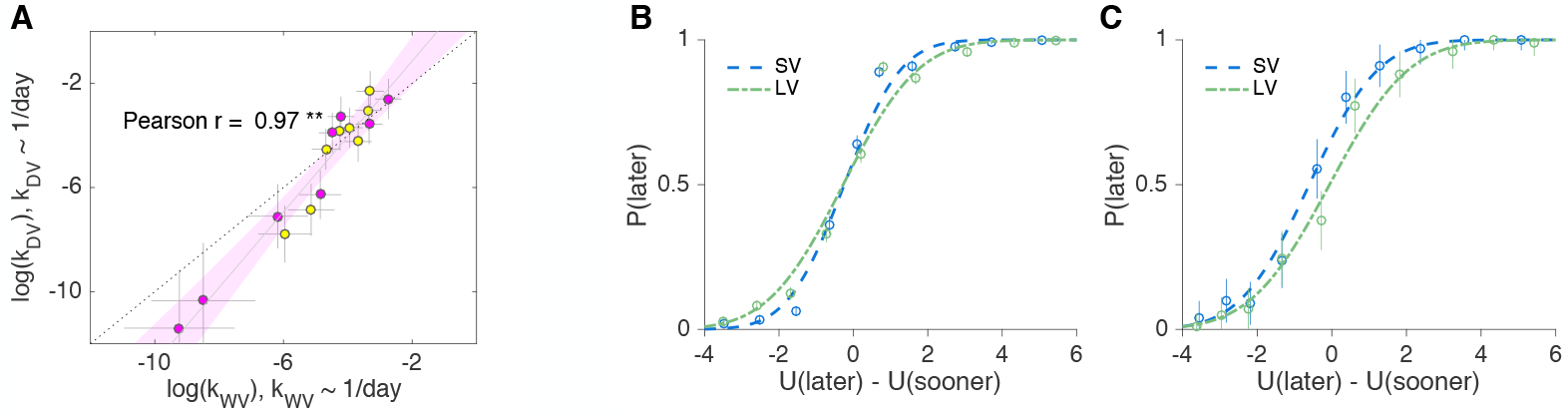
A: Control experiment 2 (n=16). The logs of delay-discounting coefficients in WV task (x-axis) plotted against the logs of delay-discounting coefficients in DV task (y-axis), *k*_*DV*_ ~ 1/*day*, *k*_*WV*_ ~ 1/*day*. The color of the circles identifies the order of task appearance. Each circle is one subject. Pearson’s r is reported on the figure. The error bars are the SD of the estimated coefficients. Solid line is the linear fit. Shaded area is the 95% CI of the linear fit. Dashed line is unity. B,C: Early trials adaptation effect. Psychometric curves for SV and LV averaged across all subjects comparing all trials (B) to first 4 trials (C).

Having ruled out the possibility that subjects ignore units of time, we test our second potential explanation: that subjects make decisions based on a subjective delay that is context dependent. We reasoned that if choices are context dependent then it may take some number of trials in each task before the context is set. Consistent with this reasoning, we found a small but significant adaptation effect in early trials: subjects are more likely to choose the later option in the first trials of SV task (Fig. 5B,C). It seems that, at first, *seconds* in the current task are interpreted as being smaller than *days* in the preceding task, but within several trials *days* are forgotten and time preferences adapt to a new time-horizon of *seconds*.

## Discussion

Using three tasks, we set out to test whether the same delay-discounting process is employed regardless of the verbal/non-verbal nature of the task and the time-horizon. We found significant correlations between subjects’ discount factors in the three tasks, providing evidence that there are common cognitive (and presumably underlying neural) mechanisms driving the decisions in the three tasks. In particular, the strong correlation between the short time-horizon non-verbal and verbal tasks (*r* = 0.79, Fig. 3A) provides the first evidence for generalizability of the non-verbal task; suggesting that this task can be applied to both human and animal research for direct comparison of cognitive and neural mechanisms underlying delay-discounting. However, the correlation between the short-delay/non-verbal task and the long-delay/verbal task is lower (*r* = 0.36). Taken together, our results suggests animal models of delay-discounting may have more in common with short time-scale consumer behavior such as impulse purchases and “paying-not-to-wait” in mobile gaming [22] and caution is warranted when reaching conclusions from the broader applicability of these models to long-time horizon real-world decisions, such as buying insurance or saving for retirement.

### Stability of preferences

The question of stability is of central importance to applying in-lab studies to real-world behavior. There are several concepts of stability that our study addresses. First, is test/re-test stability; second, stability across the verbal/non-verbal gap; third, stability across the second/day gap. Consistent with previous studies [4,40,50], we found high within-task reliability. Choices in the same task did not differ when made at the beginning or the end of the session nor when they were made in sessions held on different days even 2 weeks apart (*SI Results*).

To our knowledge, there are no studies comparing stability across the verbal/non-verbal gap for delay-discounting. The closest literature that we are aware finds that value encoding (the convexity of the utility function) but not probability weighting is similar across the verbal/non-verbal gap in sessions that compare responses to a classic verbal risky economic choice task with an equivalent task in the motor domain [81]. It may be that unlike time or value, probability is processed differently in verbal vs. non-verbal settings [33].

There are two aspects to the time-horizon gap that may contribute independently to differences in subjects’ preferences between our short and long tasks. First, there is the difference in order of magnitudes of the delays. Second, there is a difference in the experience of the delay, in that all delays are experienced in the short tasks, but only one delay is experienced in the long task.

Our control study comparing discounting of days vs. weeks eliminated the second factor since only one delay was experienced for both days and weeks tasks. We found almost perfect correspondence between the choices in the two tasks (Fig. 5A): subjects discounted 7 days as much as they discounted one week. However, days and weeks are only separated by one order of magnitude, while seconds vs. days are five orders of magnitude apart. So while the days/weeks experiment provides some evidence that the magnitude of the delays does not contribute substantially to variance in choice, it may be that larger differences (e.g. comparing hours vs. weeks) may produce an effect. The evidence from the literature on this issue is mixed. On the one hand, some have found that measures of discount factors on month long delays are not predictive of discount factors for year-long horizons (a difference of one order of magnitude) [44,74] but others have found consistent discounting for the same ranges [36]. Other studies that compared the population distributions of discount factors for short (up to 28 days) to long (years) delays (2 orders of magnitude) found no differences in subjects’ discount factors [2,21]. Some of these discrepancies can be attributed to the framing of choice options: standard larger later vs. smaller sooner compared to negative framework [44], where subjects want to be paid more if they have to worry longer about some negative events in the future.

Several previous studies have compared discounting in experienced delay tasks (as in our short tasks) with tasks where delays were hypothetical or just one was experienced [36, 40, 53, 64]. For example, Lane et al [40], also used a within-subject design to examine short vs. long delays (e.g. similar to our short-verbal and long-verbal tasks) and found similar correlations (*r* ~ 0.5 ± 0.1) with a smaller sample size (n=16). Consistent with our findings, they found (but did not discuss) a 5 order of magnitude scaling factor between subjects discounting of seconds and days suggesting that this scaling is a general phenomenon.

### Cost of waiting vs. discounting future gains

It may seem surprising that human subjects would discount later rewards, i.e. choosing immediate rewards, in a task where delays are in seconds. After all, subjects cannot consume earnings immediately. Yet, this result is consistent with earlier work that suggests individuals derive utility from receiving money irrespective of when it is consumed [48,49,63]. In our design, a pleasing (as reported by subjects) ‘slot machine’ sound accompanied the presentation of the coins in the short-delay tasks. This sound can be interpreted as an instantaneous secondary reinforcer [38]. Further, this result is consistent with studies which find that humans exhibit discount rates comparable to other species when consuming liquid rewards [35]. On the other hand, this would not be surprising for those who develop (or study) “pay-not-to-wait” video games [22], which exploit player’s impulsivity to acquire virtual goods with no actual economic value.

Using a seconds time-horizon may lead one to question if we can measure delay-discounting or if we are capturing the cost of waiting [52]. Waiting or doing nothing, “builds up anxiety and stress in an individual due both to the sense of waste and the uncertainty involved in a waiting situation” [54]. These different interpretations (Paglieri [55] described the delayed option being framed as ‘waiting’ in seconds compared to ‘postponing’ in days time-horizon) may lead one to question what we can learn from comparing within-subject behavior across tasks. Although it is not known how time is perceived, e.g. subjects could overestimate the duration of the short delay, which will lead to greater discounting, we argue that the significant correlations observed indicate there are some shared biological mechanisms underlying each of the three delay-discounting tasks, which could explain why our inability to resist a candy in a seconds time-horizon self-control task predicts our ability to complete college and other long time-horizon behaviors [13,19,28,51] (but see [77]).

### Subjective scaling of time

The range of rates of discounting we observed in the long-verbal task was consistent with that observed in other studies. For example, in a population of more than 23,000 subjects the log of the discount factors ranged from -8.75 to 1.4 ( [69], compare with Fig. 3B). This implies that, in our short tasks, subjects are discounting extremely steeply. i.e. they are discounting the rewards *per second* about the same amount that they discounted the reward *per day*. This discrepancy has been reported before [40,53]. We consider three (non-mutually exclusive) explanations for this scaling. First, subjects may ignore units. However, by testing overlapping time-horizons of days and weeks we confirmed that subjects can pay attention to units. Second, it may be that the costs of waiting [14,53,55] (discussed above) compared to the cost of postponing is, coincidentally, the same as the number of seconds in a day.

We feel this coincidence is unlikely, and thus favor the third explanation: temporal context. When making decisions about seconds, subjects ‘wait’ for seconds and when making decisions about days subjects ‘postpone reward’ for days [55]. Although our experiments were not designed to test whether the strong effect of temporal context was due to normalizing, existence of extra costs for waiting in real time, or both, we did find some evidence for the former (Fig. 5C). Consistent with this idea, several studies have found that there are both systematic and individual level biases that influence how objective time is mapped to subjective time for both short and long delays [80,83]. Thus, subjects may both normalize delays to a reference point and introduce a waiting cost at the individual level that will lead short delays to seem as costly as the long ones.

## Materials and Methods

### Participants

For the main experiment, participants were recruited from the NYU Shanghai undergraduate student population on two occasions leading to a total sample of 67 (45 female, 22 male) NYU Shanghai students. Using posted flyers, we initially recruited 35 students but added 32 more to increase statistical power (the power analysis indicates that the total of 63 participants is adequate to detect a medium to strong correlation across subjects, *SI Results*).

The study was approved by the IRB of NYU Shanghai. The subjects were between 18-23 years old, 34 subjects were Chinese Nationals (out of 67). They received a 30 CNY (~$5 USD) per hour participation fee as well as up to an additional 50 CNY (~$8 USD) per session based on their individual performance in the task (either in NV task, or total in SV and LV tasks, considering the delay of payment in the LV task). The experiment involved five sessions per subject (3 non-verbal sessions followed by 2 verbal sessions), permitting us to perform within-subject analyses. The sessions were scheduled bi-weekly and took place in the NYU Shanghai Behavioral and Experimental Economics Laboratory. In each session, all decisions involved a choice between a later (delay in seconds and days) option and an immediate (now) option. Three subjects did not pass the learning stages of the NV task. One subject did not participate in all of the sessions. These four subjects were excluded from all analyses.

### Experimental Design

The experiments were constructed to match the design of tasks used for rodent behavior in Prof. Erlich’s lab. For the temporal discounting task, the value of the later option is mapped to the frequency of pure tone (frequency ∝ reward magnitude) and the delay is mapped to the amplitude modulation (modulation period ∝ delay). The immediate option was the same on all trials for a session and was unrelated to the sound.

Through experiential learning, subjects learned the map from visual and sound attributes to values and delays. This was accomplished via 6 learning stages (0, 1, 2, 3, 4, 5) that build up to the final non-verbal task (NV) that was used to estimate subjects’ discount-factors. Briefly, the first four stages were designed to (0) learn that a mouse-click in the middle bottom ‘reward-port’ produced coins (that subjects knew would be exchanged for money), (1) learn to initiate a trial by a mouse-click in a highlighted port, (2) learn ‘fixation’: to keep the mouse-cursor in the highlighted port, (3) associate a mouse-click in the blue port with the sooner option (a reward of a fixed 4 coin magnitude that is received instantly) (4) associate varying tone frequencies with varying reward at the yellow port (5) associate varying amplitude modulation frequencies with varying delays at the yellow port. On each trial of the stage 3,4 & 5 there was either a blue port or a yellow port (but not both). The exact values for reward and delay parameters experienced in the learning stages correspond to values that are used throughout the experiment. After selecting the yellow-port (i.e. the delayed option), a countdown clock appeared on the screen and the subject had to wait for the delay which had been indicated by the amplitude modulation of the sound for that trial. Any violation (i.e. a mouse-click in an incorrect port or moving the mouse-cursor during fixation) was indicated by flashing black circles over the entire “poke” wall accompanied by an unpleasant sound (for further demonstration of the experimental time flow, please see the videos in *SI Movies*).

When a subject passed the learning stages (i.e., four successive trials without a violation in each stage, *SI Results* and Fig. S5), they progressed to the decision stages of the non-verbal task (NV). Progressing from the learning stages, a two-choice decision is present where the subject can choose between an amount now (blue choice) versus a different amount in some number of seconds (yellow choice). During the decision stages the position of blue and yellow circles on the poke wall was randomized between left and right and was always symmetrical (Fig. 1). Each of the 3 non-verbal sessions began with learning stages and continued to the decision stages. In the 2nd and the 3rd non-verbal sessions, the learning stages were shorter in duration.

The final two sessions involved verbal stimuli. During each session, subjects experience an alternating set of tasks: short delay (SV) - long delay (LV) - SV - LV (or LV-SV-LV-SV, counter-balanced per subject). An example of a trial from the short time-horizon task (SV) is shown in the sequence of screens presented in Fig. 1. The verbal task in the long time-horizon (LV) includes Initiation, Decision (as in Fig. 1) and the screen that confirms the choice. There are two differences in the implementation of these sessions relative to the non-verbal sessions. First, the actual reward magnitude and delay are written within the yellow and blue circles presented on the screen, in place of using sounds. Second, in the non-verbal and verbal short delay sessions, subjects continued to accumulate coins (following experiential learning stages) and the total earned was paid via electronic payment at the end of each experimental session. In the long-verbal sessions, a single trial was randomly selected for payment (method of payment commonly used in human studies with long delays, [17]) and shown at the end of the session. The associated payment is made now or later depending on the subject’s choice in the selected trial.

### Analysis

For model-based analysis we use hierarchical Bayesian analysis (HBA) **brms**, 2.0.1 [10,12] that allows for pooling data across subjects, recognizing individual differences and finding full posterior distributions, rather than point estimates of parameters. The means of HBA posteriors of the individual discount-factors for each task are almost identical to the individual fits done for each experimental task separately using maximum likelihood estimation through **fmincon** in Matlab (*SI Method*, Fig. S1 & Fig. S2). We further validated the HBA method by simulating choices from a population of ‘agents’ with known parameters and demonstrating that we could recover those parameters given the same number of choices per agent as in our actual dataset. The first non-verbal session data was excluded from model-fitting due to a comparatively high proportion of first-order violations than in the following two non-verbal sessions (see further discussion in *SI Results*). A 6 population level and 4 subject level parameters model (mixed-effects model (*M*_6*p*,4*s*_), *SI Method*) is used to estimate discount-factors and decision-noise from choices. At the subject level this model transforms the stimulus and individual preferences on each trial (inputs to the model include rewards and delays for sooner and later options) into a probability distribution about the subject’s choice. For the non-verbal task, we assumed that the subjects had an unbiased estimate of the meaning of the frequency and AM modulation of the sound. For example, for a given set of parameters the model would predict that trial one will result in 80% chance of the subject choosing later option. First, rewards and delays are converted in the subjective value of each choice option using hyperbolic utility model (Eq. 1). Then, Eq. S3 (a logit, or softmax function) translates the difference between the subjective value of the later and the subjective value of the sooner (estimated using Eq. 1) into a probability of later choice for each subject. Two functions below rely on the four parameters (*k*_*i,s*_: (*k*_*i,NV*_,*k*_*i,SV*_,*k*_*i,LV*_), the discounting factor per subject*task, and *τ*_*i*_ individual decision noise).

Hyperbolic utility model:

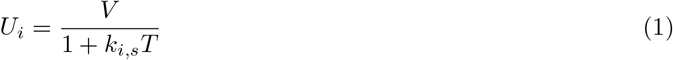

where *V* is the current value of delayed asset and *T* is the delay time.

Softmax rule:

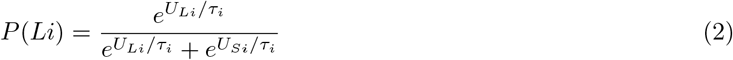

where *L* is the later, *S* is the sooner offer and *τ*_*i*_ is the individual decision noise.

For plotting posteriors of *log*(*k*) we calculated probability density estimates (for smoothing) using the **ksdensity** function in Matlab. The estimate is based on a normal kernel function, and is evaluated at equally-spaced 100 points, *x*_*i*_, that cover the range of the data in *x*.

To test for differences across tasks we examined the HBA fits using the **brms::hypothesis** function. This function allows us to directly test the posterior probability that the *log*(*k*) is shifted and/or scaled between treatments. This function returns an “evidence ratio” which tells us how much we should favor the hypothesis over the inverse (e.g. 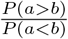) and we used Bayesian confidence intervals to set a threshold (*p* < 0.05) to assist frequentists in assessing statistical significance.

The bootstrapped (mean, median and variance) tests are done by sampling with replacement and calculating the sample statistic for each of the 10000 boots, therefore creating a distribution of bootstrap statistics and (i) testing where 0 falls in this distribution for unpaired tests or (ii) doing a permutation test to see whether the means are significantly different for paired tests.

Simulations done for both model-based and model-free analyses are described in detail in *SI Results*.

### Software

Tasks were written in Python using the **PsychoPy** toolbox (1.83.04, [58]). All analysis and statistics was performed either in Matlab (version 8.6, or higher, The Mathworks, MA), or in R (3.3.1 or higher, R Foundation for Statistical Computing, Vienna, Austria). R package **brms**(2.0.1) was used as a wrapper for Rstan [32] for Bayesian nonlinear multilevel modeling [10], **shinystan** [27] was used to diagnose and develop the brms models. Package **lme4** was used for linear mixed-effects modeling [6].

### Data Availability

Software for running the task, as well as the data and analysis code for regenerating our results are available at github.

## Acknowledgments

This work was supported by NYU Shanghai Research Challenge Fund (to S.F.L. and J.C.E.), National Science Foundation of China (NSFC-31750110461) and Shanghai Eastern Scholar Program (SESP) (to E.L.). J.C.E. was additionally supported by (1) Sponsored by Program of Shanghai Academic/Technology Research Leader (15XD1503000); (2) and by Science and Technology Commission of Shanghai Municipality (15JC1400104). We thank Paul Glimcher, Ming Hsu, & Joseph Kable for fruitful discussions. We thank NYU Shanghai undergraduate students Stephen Mathew, Xirui Zhao, Wanning Fu and Jonathan Lin who helped collect data for control experiments.

## Supporting Information

### SI Method: Functional Forms and Estimation Methods

#### Nonlinear Models

Time discounting is the decline in the subjective value of a good as a function of the expected delay and reward. There is no consensus which functional form of delay-discounting best describes human behavior. Although the exponential model [67] of time discounting has a straightforward economic meaning: a constant probability of loss of reward per waiting time, the hyperbolic model [47] seems to more accurately describe how individuals discount future rewards, in particular preference reversals [7]. In order to be sure that our results and main conclusions did not depend on the method (e.g. hierarchical Bayesian vs. maximum likelihood estimation of individual subject parameters) or functional form (e.g. exponential vs. hyperbolic), we validated our results with several methods.

Exponential model:

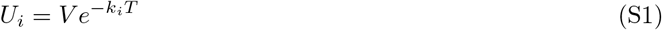

Hyperbolic model:

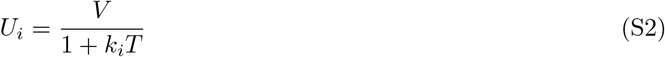

where *V* is current value of delayed asset, *T* is the delay time and *k_i_* is the individual discounting factor.

We considered both a shift-invariant softmax rule and a scale-invariant matching rule to transform the subjective utilities of the sooner and later offers into a probability of choosing the later offer.

Softmax:

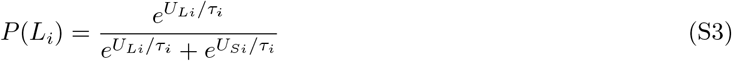

Matching rule:

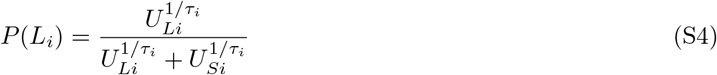

where *L*_*i*_ is the later, *S*_*i*_, is the sooner option, and *τ*_*i*_ is the individual decision rule noise, or temperature.

Using maximum likelihood estimation we fit each subject’s choices to four baseline models: 1) hyperbolic utility with softmax, 2) exponential utility with softmax, 3) hyperbolic utility with matching rule and 4) exponential utility with matching rule. We also considered models that account for utility curvature, i.e. *V* is replaced by 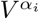 and models that account for trial number and cumulative waiting time.

In the models that account for trial number or cumulative wait time the individual discounting factor, *k*_*i*_ consists of a constant component *k*_*ci*_ and time-dependent component:

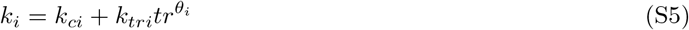

or

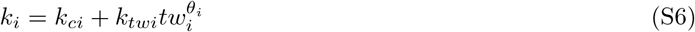

where *tr* is the trial number and *tw*_*i*_ is the individual’s total waiting time in seconds that exists only for short delays task and *θ*_*i*_ is a scaling parameter.

#### MLE Estimation Methods

We first estimated subjects’ time-preferences individually (since discounting factors differ among people) for each experimental task with maximum likelihood estimation (MLE) and used leave-one-trial-out cross-validation for model comparison. Based on the Bayesian information criterion criterion (BIC, Table S1) and number of subjects that were well described by the model (Table S2), the softmax-hyperbolic model was selected as the best nonlinear model, and we used this for the Bayesian Hierarchical modeling.

**Table S1.** BIC Comparison for Various Models (the smaller the better)

|  | BIC | SE |
| --- | --- | --- |
| Matchrule-Hyperbolic | 179.47 | 4.99 |
| Softmax-Hyperbolic | 191.16 | 4.96 |
| Matchrule-Exponential | 192.03 | 4.82 |
| Softmax-Hyperbolic-Time-Trial | 210.00 | 4.77 |
| Softmax-Hyperbolic-Time-Wait | 213.23 | 4.98 |
| Softmax-Exponential | 215.33 | 4.45 |
| Softmax-Hyperbolic-Curvature | 222.98 | 4.18 |

Since the estimation procedure was identical for all MLE fits (Matlab code available on github repository) we describe it using the softmax-hyperbolic model as an example. This is a 2-parameter model to estimate choice behavior, i.e. it transforms the stimulus on each trial (inputs to the model include rewards and delays for sooner and later options) into a probability distribution about the subject’s choice. For example, if for a given set of parameters, the model predicts that trial 1 will result in 80% chance of the subject choosing later option, and the subject, in fact, chose the later, the trial would be assigned a likelihood of 0.8 (if the subject chose sooner, the trial would have a likelihood of 0.2). Finally, we perform a leave-one-out cross-validation for each subject-task to avoid overfitting. We leave one trial out and use the rest of the trials in the experimental task to predict this trial. We repeat this procedure for each trial.

Figure S1 shows an example of the model fit and estimated parameters for one of the subjects. For this particular subject fitting softmax-hyperbolic model in the non-verbal task resulted in delay-discounting coefficient *k* = 0.07 (Fig. S1). We can readily observe that although first-order violations were present during non-verbal task, in the verbal task they get eliminated. Although the unit-free discount rates seem pretty stable, the decision noise *τ*_*i*_ gets smaller from non-verbal to verbal tasks.

**Figure S1.**
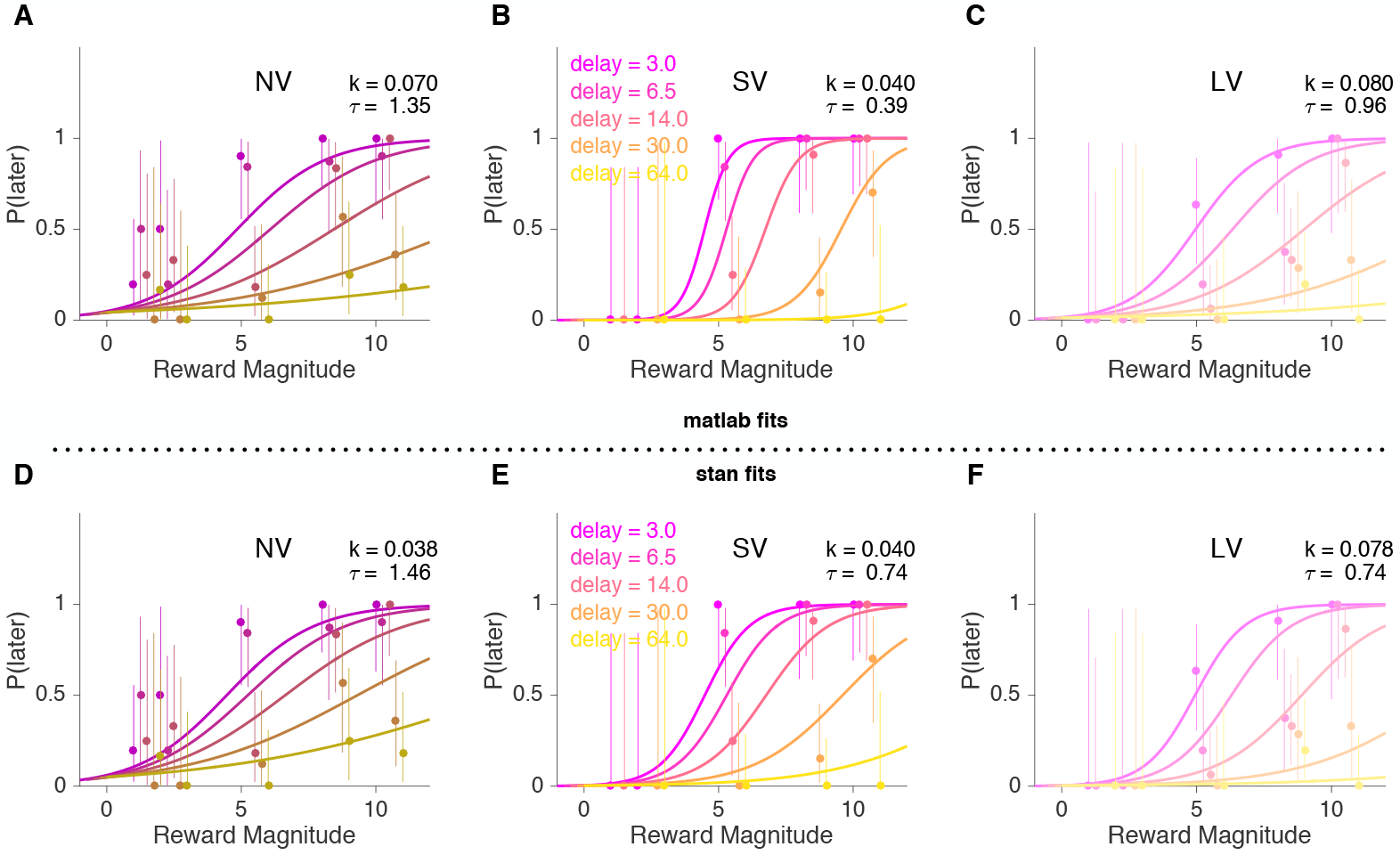
An example of the softmax-hyperbolic fit for one subject. A-C - Matlab fits, D-F - Stan fits. Psychometric curves obtained from the model fits versus actual data (circles) for non-verbal (NV) and verbal (short (SV) and long (LV) delay) tasks for each delay value, where the x-axis is the reward and the y-axis is the probability (or proportion for actual choices) of later choice. Error bars are binomial.

**Table S2.** Top 3 Models Rank Correlations

| | Model | $N$ | NV vs. SV | SV vs. LV |
| --- | --- | --- | --- | --- |
| $\log(k_i)$ | M-Hyperbolic | 57 | 0.52, $p < 0.01$ | 0.46, $p < 0.01$ |
| $\log(k_i)$ | S-Hyperbolic | 63 | 0.68, $p < 0.01$ | 0.49, $p < 0.01$ |
| $\log(k_i)$ | M-Exponential | 49 | 0.38, $p < 0.01$ | 0.5, $p < 0.01$ |

**Figure S2.**
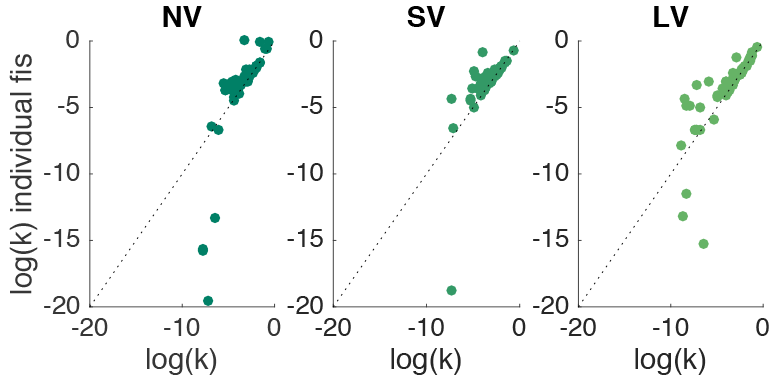
HBA model pooled data Stan fits (x-axis) vs. Matlab individual data fits (y-axis) across tasks against the unity line.

#### Hierarchical Bayesian Estimation Methods

Instead of getting point estimates (or distributions of parameter values through cross-validation as we did) one can use hierarchical Bayesian analysis (HBA) to find full posterior distributions. It allows for both pooling data across subjects and recognizing individual differences. After comparing MLE fits we decided to use the softmax-hyperbolic functional form as described above.

Using the **brms** [10] package in R allows to do HBA of nonlinear multilevel models in Stan [12] with the standard R formula syntax:

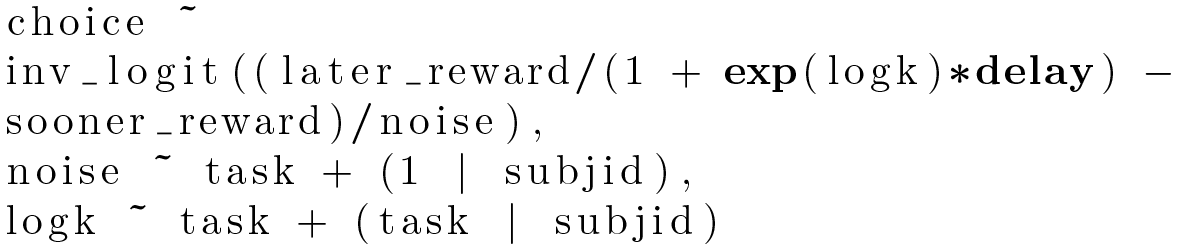

where later_reward is the later reward, sooner_reward is the sooner reward; logk is the natural logarithm of the discounting parameter *k* and noise (*τ*;) is the decision noise (like in Eq. S2 and S3, respectively). The model (*M*_6*p*,4*s*_) had 6 population level parameters (discount factor, *k*, and decision-noise, *τ* for each of the three tasks) and 4 parameters per subject: *k*_*NV*_,*k*_*SV*_,*k*_*LV*_ and *τ*. The parameters from this model are used in the main text and in SI Results.

### SI Results

#### Significant Rank Correlations Are Robust to Different Functional Forms and Methods of Estimation

Table S2 combines rank correlations obtained from individual level MLE fits from top 3 models by BIC: ‘model’ - functional form that is fitted (‘M-’ is matchrule, ‘S-’ is softmax, models are presented in the order of BIC), ‘*N*’ - number of models converged out of 63 (total number of subjects), finally, Spearman *r* for rank correlations between non-verbal and short delay verbal tasks (NV vs. SV) and short delay verbal vs. long delay verbal tasks (SV vs. LV), respectively.

The fits from HBA model are almost identical to the individual fits done for each experimental task separately using softmax-hyperbolic model and ‘fmincon’ function in Matlab (Fig. S2). The individual *log*(*k*) values also agree with the range of delay-discounting values reported in a large cohort GWAS delay-discounting study [69]. The rank correlation values for individual fits in table S2 correspond to the ones in the main text both in magnitude and significance.

For nonparametric analysis we calculated a coarse model-free measure: percentage of trials in which the later option was chosen in each of the three tasks (percent ‘yellow’ choice). There was significant heterogeneity between subject’s responses (NV: mean = 0.45, std = 0.19; SV: mean = 0.59, std = 0.23; LV: mean = 0.59, std = 0.26; our experiments considered both larger and smaller later options, with the percentage of smaller later options bigger for nonverbal than for verbal tasks, see further discussion on first-order violations). Nevertheless, if we ranked the subjects by the fraction of trials they chose the later option in each task, there is a strong correlation between subjects’ ranks across tasks (SV vs. NV, Spearman *r* = 0.71 (*p* < 0.01); SV vs. LV, *r* = 0.49 (*p* < 0.01); NV vs. LV, *r* = 0.30 (*p* < 0.05), regardless of gender or nationality (see ‘Significance Tests of Demographic and Psychological Categories’ below).

#### First-order Violations

The notion of first-order stochastic dominance is usually defined for gambles [75,82]. Compliance with first-order stochastic dominance means that, in principle, this observed behavior can be adequately modeled with a utility-function style analysis. Utility-function analysis is controversial in part if subjects show inconsistent behavior and require significant variation within subjects. By design, our experiments considered both larger and smaller later options generating significant variation to identify utility function parameters. Overall, in the non-verbal task there was 25% of smaller later options, whereas in the verbal experiment there was 10% of smaller later options. Given that the smaller later option is always strictly worse than larger immediate option, if in such a trial smaller later is chosen, economic theory would classify this choice as reflecting a first-order violation. In the non-verbal task, violations could result from lapses in attention, motor errors or difficulty in transforming the perceptual stimuli into offers (in particular, early on in the first session while learning has not completed). In the verbal tasks, inattention and/or misunderstanding are likely explanations of violations. Our analysis indicates that first-order violations declined significantly from non-verbal (16%) to verbal tasks (4%) (Wilcoxon signed-rank test, *p* < 0.01) and are not dependent on gender or nationality. The proportion of first-order violations decreased from 26% of trials in the first non-verbal session (*NV*_1_) to 19% and 13% for the next two non-verbal sessions, *NV*_2_ and *NV*_3_, respectively (Wilcoxon signed-rank test, *NV*_1_ vs. *NV*_2_ & *NV*_1_ vs. *NV*_3_, *p* < 0.01). It is important that *NV*_1_ did not differ significantly in choice consistency (the number of preference reversals was not significantly different between *NV*_1_ and later non-verbal sessions, Wilcoxon signed-rank test, all *p* > 0.2).

#### Significance Tests of Demographic and Psychological Categories

We don’t find any significant effects for any of the categorical subjects’ groups, including gender and nationality in learning stages (*Learning Stages Analysis*), intertemporal decisions and first-order violations. For the proportion of ‘yellow’ choice there is no significant difference between females and males (females: mean = 0.5636, std = 0.2429; males: mean = 0.5322, std = 0.2900; Wilcoxon rank sum test, *p* = 0.22) and between Chinese and Non-Chinese (Chinese: mean = 0.5666, std = 0.2341; Non-Chinese: mean = 0.5384, std = 0.2851; Wilcoxon rank sum test, *p* = 0.3309). Similarly, for the first-order violations there is no significant difference between females and males (females: mean = 1.1353 (violations per session), std = 1.9655; males: mean = 1.0682, std = 2.1959; Wilcoxon rank sum test, *p* = 0.2607) and a slight difference between Chinese and Non-Chinese (Chinese: mean = 1.2086, std = 2.0684; Non-Chinese: mean = 1.0031, std = 2.0039; Wilcoxon rank sum test, *p* < 0.1).

We used the Barratt Impulsiveness Scale (BIS-11; [57]) as a standard measure of impulsivity. This test is reported to often correlate with biological, psychological, and behavioral characteristics. The mean total score for our students sample was 61.79 (std = 9.53), which is consistent with other reports in the literature (e.g., [71]). The BIS-11 did not correlate significantly with the estimated delay-discounting coefficients (BIS vs. *log*(*k*_*NV*_): Pearson *r* = 0.2, *p* = 0.1180; BIS vs. *log*(*k*_*SV*_): Pearson *r* = 0.19, *p* = 0.1384; BIS vs. *log*(*k*_*LV*_)): Pearson *r* = 0.15, *p* = 0.2521).

**Figure S3.**
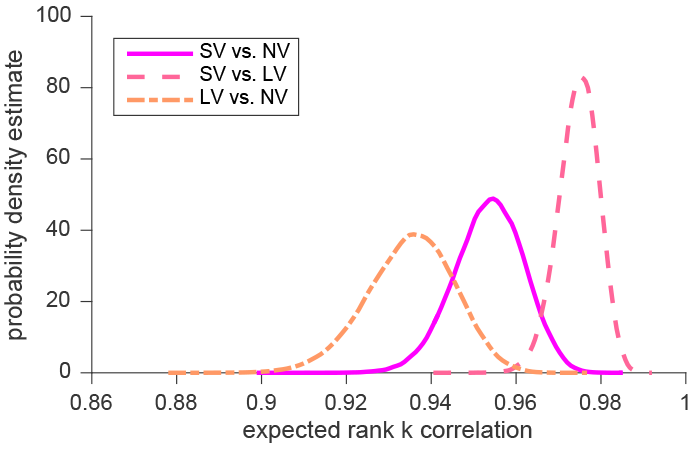
Simulation results: distributions of expected correlations of the delay-discounting coefficients ranks between tasks.

**Table S3.** Corrected Correlations of subjects’ discount factors

|  | Spearman<br>Rank Correlation | Corrected<br>Rank Correlation |
| --- | --- | --- |
| SV vs. NV | 0.76 | 0.77 |
| SV vs. LV | 0.54 | 0.57 |
| NV vs. LV | 0.36 | 0.39 |
| all $p < 0.01$ | | |

#### Model-based Analysis (Simulations)

Given that we are estimating subjects discount-factors using a finite number of trials per task, even if subjects’ discount-factors were identical in different tasks, we would not expect the rank correlations to be perfect. In order to estimate the expected maximum correlation we could observe we simulated ‘consistent” subjects’ choices using hyperbolic model with softmax rule, assuming that there is a single delay-discounting parameter - *k*_*i*_ (mean across tasks) for each subject. We did this 100,000 times and computed a distribution of pairwise rank correlations (Fig. S3). These simulations revealed that the decision noise contributed a very small amount of variance (Table S3).

#### Model-free Analysis (Nonparametric Predictions)

To validate our model-free analysis we did a non-parametric out-of-sample prediction by binning each subject’s choices into 7 bins (7-vectors, based on delay & reward) for each task (21 bins total). A 7-vector is a way of clustering data into 1) ‘small’ rewards (1 or 2 coins), 2) ‘medium’ reward and ‘small’ delay (5 coins in 3 or 6.5 secs/days), 3) ‘medium’ reward and ‘medium’ delay (5 coins in 14 secs/days), 4) ‘medium’ reward and ‘large’ delay (5 coins in 30 or 64 secs/days), 5) ‘large’ reward and ‘small’ delay (8 or 10 coins in 3 or 6.5 secs/days), 6) ‘large’ reward and ‘medium’ delay (8 or 10 coins in 14 secs/days), and 7) ‘large’ reward and ‘large’ delay (8 or 10 coins in 30 or 64 secs/days).

Next, we combined the 7-vectors for each subject into a 21-vector and performed a leave-one-subject/task-out cross-validation. For each subject-task, we trained a linear model to predict the choices from one task based on the other two tasks for all other subjects. Then, for the left out subject we predicted each task from the other two. We can predict 70% of the subjects 21-vectors this way (Fig. S4). We conclude that, for most subjects, there is a shared scaling effect that allows each tasks’ choices to be predicted by the other two.

#### Time and Reward Re-Scaling

In our main experimental tasks we used two types of delays: in seconds and in days, where 1 day = 86400 seconds. We also used three types of exchange rates: for non-verbal task 1 coin = 0.1 RMB; for verbal short delay 1 coin = 0.05 RMB; for long delay 1 coin = 4 RMB. Humans tend to discount large rewards less steeply than small rewards, i.e. discounting rates tend to increase as amounts decrease [29,30]. We re-calculated the model-based (softmax-hyperbolic model) median HBA model fits: 1) we convert them to the same units (1/days): *log*(*k*_*NV*_) = 4173.1 (by multiplying original unit-free *k* by the day to seconds conversion rate), *log*(*k*_*SV*_) = 2548.8, *k*_*LV*_ = 0.0356, 2) we consider reward re-scaling: “going from $10 to $.20, a factor of 50, k values would increase by a factor of 2” [53] *k*_*NV*_ = 4173.1, *k*_*SV*_ = 2548.8, *k*_*LV*_ = 0.0712 and 3) conclude that discrepancy of discount rates between time-horizons cannot be accounted by magnitude effects. Thus, the discount rate revealed in the verbal short delay task is more than 10^4^ times larger than the rate describing the choices made by the same participants in the verbal long delay task.

**Figure S4.**
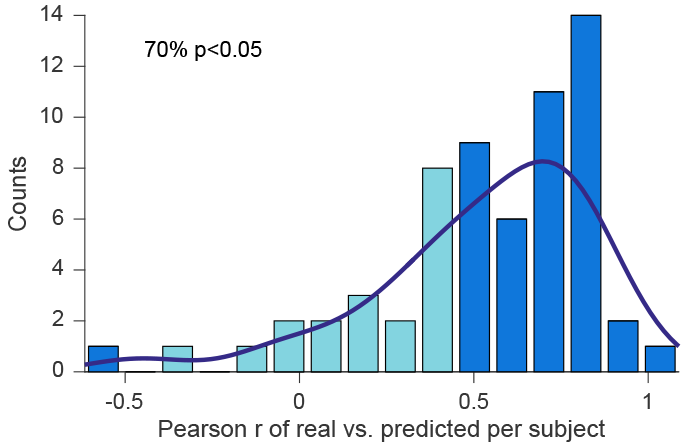
Distribution of the Pearson correlation coefficients between real and predicted 21-vectors of each subject. Darker bars are significant correlations.

#### Dynamic Stability across and within Sessions

We checked for possible dynamic instability within experimental tasks. We find that time-preferences (measured as percent ‘yellow’ choices) are stable in the same task between the first half of the block and the second half of the block for each subject (Wilcoxon signed-rank test, *p* = 0.3491). Similar to other researchers [4, 50] we observe stability within subjects across experimental sessions that take place each other week: no significant difference in percent ‘yellow’ choice between NV sessions (Wilcoxon signed-rank test, *p* = 0.4721), between SV sessions (Wilcoxon signed-rank test, *p* = 0.6613) and a slight difference between LV sessions (Wilcoxon signed-rank test, *p* < 0.1).

#### Power Analysis

We ran power analysis to find out total sample size required to determine whether a correlation coefficient differs from zero. For expected correlation *r* = 0.5 and 80% power (the ability of a test to detect an effect, if the effect actually exists, [9,16]) the required sample size is N = 29, for a medium size correlation of *r* = 0.3 - N = 84.

#### Learning Stages Analysis

64 (1 out of 64 did not complete all sessions of the study, the analysis below is done for 63 subjects) out of 67 subjects passed learning stages (*SI Movies*) for our novel language-free task.

There were 6 learning stages (0, 1, 2, 3, 4, 5). The first four stages were respectively designed to 0) learn the reward port, 1) learn the initiation port, 2) fixation, and 3) associate the blue colored port with the sooner option (a reward of a fixed 4 coin magnitude that is received instantly). 4 trials without violations in a row are required to pass these learning stages. In stage 4, subjects are primed to the sound frequency to learn the variability of reward magnitudes (1, 2, 5, 8, or 10 coins): first, the lower and upper bounds, then, in ascending and descending order and, finally, in random order. In the final stage 5, subjects heard the AM of a sound during fixation that is now mapped to the delay (3, 6.5, 14, 30, or 64 seconds) of the later option. The order of the stimuli presented was the same as in the previous stage. For the last two stages 4 trials without violations in a row are required during the random order of stimuli presentation to pass. Learning stages were shorter for the 2nd and the 3rd non-verbal sessions, since only random rewards and delays were presented for ‘forced’ choices excluding the experience of the bounds and the ordered values.

**Table S4.** Percent of Trials without Violation in Each Learning Stage

|  | stage 0 | stage 1 | stage 2 | stage 3 | stage 4 | stage 5 |
| --- | --- | --- | --- | --- | --- | --- |
| mean | 0.9092 | 0.9126 | 0.3304 | 0.8482 | 0.81 | 0.7734 |
| std | 0.1567 | 0.1098 | 0.2490 | 0.16 | 0.1851 | 0.2036 |

**Figure S5.**
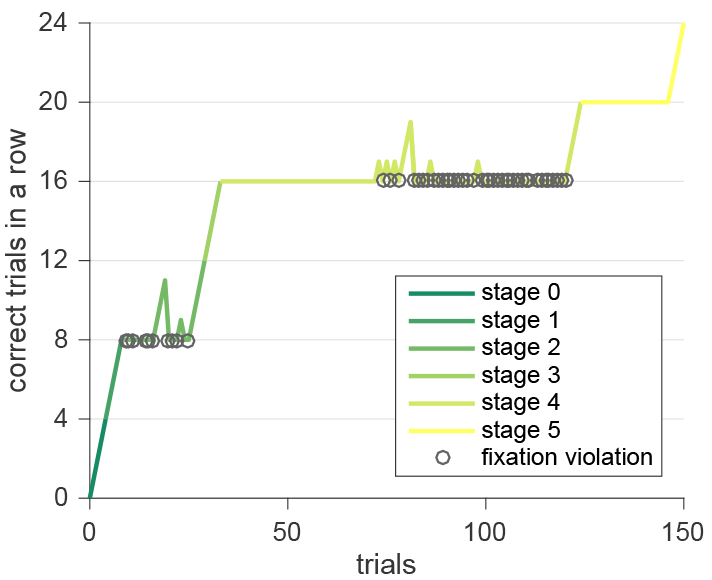
Violation Pattern 2. Each one unit increase in y with increase in x (trials) is a correct trial. Each violation drops the ‘correct in a row’ measure to 0 for each stage. 4 correct trials in a row are required to advance to the next learning stage. In stages 4 and 5 only trials with random order of stimuli were counted as correct or not correct. The circles identify fixation violations.

There were no significant differences between learning stage performance across demographic categories, such as gender and nationality (between females (mean = 0.7503 (percent trials without violations), std = 0.2814) and males (mean = 0.7874, std = 0.2713): Wilcoxon rank sum test, *p* < 0.1; between Chinese (mean = 0.7463, std = 0.2870) and Non-Chinese (mean = 0.7877, std = 0.2643): Wilcoxon rank sum test, *p* = 0.1268). Subjects experienced difficulty with the learning stage 2. This is the only stage where the average performance was less than 40% compared to more than 70% performance in other stages (Table S4). Subjects on average also spent significantly more time measured in seconds for the learning stage 2 compared to the next learning stage 3 (Wilcoxon signed-rank test, *p* < 0.01), although only 4 trials without violations are required to pass this learning stage and learning stage 3 includes more steps. During learning stage 2 subjects had to learn fixation. Fixation was specifically designed to drive subjects attention away from the computer mouse (since no movement is allowed outside of the circle during fixation) and bring focus to other senses. During fixation in the learning stages 4 and 5 (as well as in the decision stages) subjects hear sound that corresponds to the reward magnitude and delay.

There were three major patterns of violations: 1. subject had difficulty passing stage 2 (because of fixation violations), however later stages were completed quickly; 2. subject was able to pass stage 2, by having 4 correct answers in a row, but during stage 4 encountered problems with fixation violation again; 3. subject was able to proceed till stage 5 almost without violations, but was stopped by several fixation violations at stage 5. Figure S5 shows the second pattern of correct choices vs. violations.

### SI Method: Control Experiments

#### Control Experiment 1: No Circles (NC)

In total, 25 (29 started, 4 withdrew) undergraduate students from NYU Shanghai participated in 5 experimental sessions (3 non-verbal and 2 verbal sessions, in this sequence, that were scheduled bi-weekly). The study requirements in order to meet the IRB protocol conditions remained the same as in the main experiment (*Materials and Methods*). In each session, subjects completed a series of intertemporal choices. Across sessions, 160 trials were conducted in each of the following tasks mimicking the main experiment, i) non-verbal (NV), ii) verbal short delay (SV; 3 seconds - 64 seconds), and iii) verbal long delay (LV; 3 days - 64 days). In each trial, irrespective of the task, subjects made a decision between the sooner and the later options. The NV task was exactly the same as in the main experiment. All subjects passed learning stages. The SV and LV tasks differed from the main experiment in exactly two ways: 1) the stimuli presentation didn’t include a display of circles of different colors. Instead, two choices were presented on the left or on the right side (counterbalanced) of the screen (Fig. S6), 2) the subjects didn’t have to click on the circles using mouse, instead they used a keyboard to indicate ‘L’ or ‘R’ choice. Everything else stayed the same as in the main experiment, i.e the last two sessions included an alternating set of verbal tasks: SV-LV-SV-LV (or LV-SV-LV-SV, for a random half of subjects), the payment was done differently for SV and LV (randomly picked trial for payment in LV, *Materials and Methods*), etc. The purpose of this control experiment is to confirm that significant correlation between non-verbal tasks and verbal tasks we report in Results is not an artifact of our main experimental design: subjects experience the same visual display and motor responses in the non-verbal and verbal tasks and this design similarity might drive the correlation between time-preferences in these tasks. Instead, in this control experiment the verbal tasks are made as similar as possible (keeping our experiment structure) to typical intertemporal choice tasks used in human subjects.

**Figure S6.**
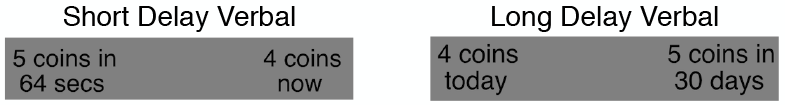
Control Experiment 1 choice screen example.

#### Control Experiment 2: Days & Weeks (DW)

In total, 16 subjects took part in this experiment. Subjects were undergraduate students from NYU Shanghai. This experiment was approved under the same IRB protocol as the control experiment 1 and the main experiment. This experiment included two following experimental tasks: i) verbal days delay (DV; 1 day - 64 days) and ii) verbal weeks delay (WV; 1 week - 35 weeks). Subjects underwent only one session where the verbal tasks were alternated: DV-WV-DV-WV (or WV-DV-WV-DV, for roughly half of subjects; 200 trials per task). For each of the tasks in this control experiment the stimuli and procedures were exactly the same as for LV task in the control experiment 1. The purpose of this control task is to check whether subjects pay attention to units.

### SI Results: Control Experiments

Individual delay-discounting fits were estimated using HBA softmax-hyperbolic model using the same procedure as in the main experiment. No-Cirles (NC) data was analyzed as is, keeping delay units in seconds and in days, whereas days-weeks (DW) data was analyzed after converting delays in weeks to days. Among estimated delay-discounting coefficients (Fig. 4C), there is no significant difference in means of *log*(*k*) between tasks (bootstrapped mean (and median) tests, SV vs. LV and NV vs. SV, all *p* > 0.8). Thus, we find that similar to the main experiment there is no common shift across tasks and individual effects per task explain a significant amount of variance. For DW, based on Fig. 5A we conclude that day delay task and week delay task are likely to be perceived by subjects as a single delay task with different units in it. Subjects do pay attention to units and individual differences between delay-discounting coefficients do matter, while the differences between the tasks do not.

#### Correlations Between Tasks and Order Effects

Even with a smaller subject’s pool (25 subjects) for the NC control experiment the correlation of ranks of *log*(*k*) between SV and NV tasks stays strong, while the Pearson correlation becomes a bit smaller (Fig. 4A,B; Table S5). To determine whether the correlations observed were within the range expected by chance, we repeatedly (10,000 times) randomly sampled 25 of the original 63 subjects (from Fig.) and computed the correlations between tasks. Pearson *r* = .42 is lower than we would expect for NC (the 95% CI of the correlation assuming 25 subjects is [0.498 0.915]). This suggests that some of the SV-NV correlation in the main experiment may be driven by visuo-motor similarity in experimental designs.

**Table S5.** Correlations of Subjects’ Discount-factors for NC

|  | Pearson<br>Correlation | Spearman<br>Rank Correlation |
| --- | --- | --- |
| SV vs. NV | 0.42* | 0.52** |
| SV vs. LV | 0.69** | 0.52** |
| NV vs. LV | 0.36 | 0.33 |

\*\* $p < 0.01$ , \* $p < 0.05$

The correlation between *log*(*k*) and ranks of *log*(*k*) for DW experiment is almost perfect (Fig. 4D; Pearson *r* = .97; Spearman *r* = .95, all *p* < 0.01). This suggests that subjects are making choices in day and week delay tasks by converting these delays to a common unit.

We don’t find any order effects for the NC control experiment (bootstrapped mean test, SV-LV-SV-LV vs. LV-SV-LV-SV order for SV and LV *log*(*k*), respectively, all *p* > 0.6) as well as for the DW control experiment (bootstrapped mean test, DV-WV-DV-WV vs. WV-DV-WV-DV order for DV and WV *log*(*k*), respectively, all *p* > 0.2). To confirm the absence of order effect we also run an order model with DW data, where (*log*(*k*) ~ (*order*|*subjid*)). The comparison based on 10-fold cross validation criteria (using the ‘kfold’ function in the ‘brms’ package) between order and main models is in favor of the latter (order = 3467.59 ¿ 3261.15 = main), since lower is better.

### SI Movies: Experimental Task

#### Learning

We provide videos of the learning stages, showing the examples of violations that can be made. The video starts with stage 0 - 00:00 and continues with stage 1 - 00:14, stage 2 - 00:31, stage 3 - 00:55, stage 4 (trimmed) - 01:18 and stage 5 (trimmed) - 01:41.

#### NV, SV, LV

In addition, videos of several trials of decision stages for non-verbal, short delay and long delay tasks are recorded.

## References

1. D. Åkerlund, B. H. Golsteyn, H. Grönqvist, and L. Lindahl. Time discounting and criminal behavior. Proceedings of the National Academy of Sciences, page 201522445, 2016.

2. S. Andersen, G. W. Harrison, M. I. Lau, and E. E. Rutström. Discounting behavior: A reconsideration. European Economic Review, 71:15–33, 2014.

3. J. Andreoni, M. A. Kuhn, and C. Sprenger. Measuring time preferences: A comparison of experimental methods. Journal of Economic Behavior & Organization, 116:451–464, 2015.

4. N. Augenblick, M. Niederle, and C. Sprenger. Working over time: Dynamic inconsistency in real effort tasks. The Quarterly Journal of Economics, page qjv020, 2015.

5. D. Bates, M. Mächler, B. Bolker, and S. Walker. Fitting linear mixed-effects models using lme4. arXiv preprint arXiv:1406.5823, 2014.

6. D. Bates, M. Maechler, B. Bolker, S. Walker, et al. lme4: Linear mixed-effects models using eigen and s4. R package version, 1(7):1–23, 2014.

7. G. S. Berns, D. Laibson, and G. Loewenstein. Intertemporal choice-toward an integrative framework. Trends in cognitive sciences, 11(11):482–488, 2007.

8. T. C. Blanchard, J. M. Pearson, and B. Y. Hayden. Postreward delays and systematic biases in measures of animal temporal discounting. Proceedings of the National Academy of Sciences, 110(38):15491–15496, 2013.

9. D. G. Bonett and T. A. Wright. Sample size requirements for estimating pearson, kendall and spearman correlations. Psychometrika, 65(1):23–28, 2000.

10. P.-C. Bürkner. brms: An R package for bayesian multilevel models using Stan. Journal of Statistical Software, 80(1):1–28, 2017.

11. X. Cai, S. Kim, and D. Lee. Heterogeneous coding of temporally discounted values in the dorsal and ventral striatum during intertemporal choice. Neuron, 69(1):170–182, 2011.

12. B. Carpenter, A. Gelman, M. Hoffman, D. Lee, B. Goodrich, M. Betancourt, M. A. Brubaker, J. Guo, P. Li, and A. Riddell. Stan: A probabilistic programming language. Journal of Statistical Software, 20:1–37, 2016.

13. B. Casey, L. H. Somerville, I. H. Gotlib, O. Ayduk, N. T. Franklin, M. K. Askren, J. Jonides, M. G. Berman, N. L. Wilson, T. Teslovich, et al. Behavioral and neural correlates of delay of gratification 40 years later. Proceedings of the National Academy of Sciences, 108(36):14998–15003, 2011.

14. G. B. Chapman. Time preferences for the very long term. Acta psychologica, 108(2):95–116, 2001.

15. G. B. Chapman and A. S. Elstein. Valuing the future: Temporal discounting of health and money. Medical decision making, 15(4):373–386, 1995.

16. J. Cohen. Statistical power analysis for the behavioral sciences. hilsdale. NJ: Lawrence Earlbaum Associates, 2, 1988.

17. K. M. Cox and J. W. Kable. Bold subjective value signals exhibit robust range adaptation. Journal of Neuroscience, 34(49):16533–16543, 2014.

18. J. W. Dalley, B. J. Everitt, and T. W. Robbins. Impulsivity, compulsivity, and top-down cognitive control. Neuron, 69(4):680–694, 2011.

19. A. L. Duckworth, E. Tsukayama, and T. A. Kirby. Is it really self-control? examining the predictive power of the delay of gratification task. Personality and Social Psychology Bulletin, 39(7):843–855, 2013.

20. S. Dunne, A. D’Souza, and J. P. O’Doherty. The involvement of model-based but not model-free learning signals during observational reward learning in the absence of choice. Journal of neurophysiology, 115(6):3195–3203, 2016.

21. C. Eckel, C. Johnson, and C. Montmarquette. Saving decisions of the working poor: Short-and long-term horizons. In Field experiments in economics, pages 219–260. Emerald Group Publishing Limited, 2005.

22. E. Evans. The economics of free: Freemium games, branding and the impatience economy. Convergence, 22(6):563–580, 2016.

23. N. A. Fineberg, M. N. Potenza, S. R. Chamberlain, H. A. Berlin, L. Menzies, A. Bechara, B. J. Sahakian, T. W. Robbins, E. T. Bullmore, and E. Hollander. Probing compulsive and impulsive behaviors, from animal models to endophenotypes: a narrative review. Neuropsychopharmacology, 35(3):591, 2010.

24. S. Frederick, G. Loewenstein, and T. O’donoghue. Time discounting and time preference: A critical review. Journal of economic literature, 40(2):351–401, 2002.

25. B. Fung, C. Murawski, and S. Bode. Caloric primary rewards systematically alter time perception. 2017.

26. E. E. Furlong and J. E. Opfer. Cognitive constraints on how economic rewards affect cooperation. Psychological Science, 20(1):11–16, 2009.

27. J. Gabry. Shinystan: interactive visual and numerical diagnostics and posterior analysis for bayesian models. R Package Version, 2(0), 2015.

28. B. H. Golsteyn, H. Grönqvist, and L. Lindahl. Adolescent time preferences predict lifetime outcomes. The Economic Journal, 124(580):F739–F761, 2014.

29. L. Green, J. Myerson, D. D. Holt, J. R. Slevin, and S. J. Estle. Discounting of delayed food rewards in pigeons and rats: is there a magnitude effect? Journal of the experimental analysis of behavior, 81(1):39–50, 2004.

30. L. Green, J. Myerson, and P. Ostaszewski. Amount of reward has opposite effects on the discounting of delayed and probabilistic outcomes. Journal of Experimental Psychology-Learning Memory and Cognition, 25(2):418–427, 1999.

31. L. Gregorios-Pippas, P. N. Tobler, and W. Schultz. Short-term temporal discounting of reward value in human ventral striatum. Journal of Neurophysiology, 101(3):1507–1523, 2009.

32. J. Guo, D. Lee, K. Sakrejda, J. Gabry, B. Goodrich, J. De Guzman, E. Niebler, T. Heller, and J. Fletcher. rstan: R interface to stan. R, 534:0–3, 2016.

33. R. Hertwig and I. Erev. The description-experience gap in risky choice. Trends in cognitive sciences, 13(12):517–523, 2009.

34. J. Huang, Z. Zhong, M. Wang, X. Chen, Y. Tan, S. Zhang, W. He, X. He, G. Huang, H. Lu, et al. Circadian modulation of dopamine levels and dopaminergic neuron development contributes to attention deficiency and hyperactive behavior. Journal of Neuroscience, 35(6):2572–2587, 2015.

35. K. Jimura, J. Myerson, J. Hilgard, T. S. Braver, and L. Green. Are people really more patient than other animals? evidence from human discounting of real liquid rewards. Psychonomic Bulletin & Review, 16(6):1071–1075, 2009.

36. M. W. Johnson and W. K. Bickel. Within-subject comparison of real and hypothetical money rewards in delay discounting. Journal of the experimental analysis of behavior, 77(2):129–146, 2002.

37. J. W. Kable and P. W. Glimcher. The neural correlates of subjective value during intertemporal choice. Nature neuroscience, 10(12):1625–1633, 2007.

38. R. T. Kelleher and L. R. Gollub. A review of positive conditioned reinforcement. Journal of the Experimental Analysis of behavior, 5(S4):543–597, 1962.

39. M. W. Khaw, P. W. Glimcher, and K. Louie. Normalized value coding explains dynamic adaptation in the human valuation process. Proceedings of the National Academy of Sciences, page 201715293, 2017.

40. S. D. Lane, D. R. Cherek, C. J. Pietras, and O. V. Tcheremissine. Measurement of delay discounting using trial-by-trial consequences. Behavioural processes, 64(3):287–303, 2003.

41. B. Lau and P. W. Glimcher. Dynamic response-by-response models of matching behavior in rhesus monkeys. Journal of the experimental analysis of behavior, 84(3):555–579, 2005.

42. C.-T. Law and J. I. Gold. Reinforcement learning can account for associative and perceptual learning on a visual-decision task. Nature neuroscience, 12(5):655–663, 2009.

43. F. Lieder, T. L. Griffiths, and M. Hsu. Overrepresentation of extreme events in decision making reflects rational use of cognitive resources. Psychological review, 125(1):1, 2018.

44. G. Loewenstein and R. H. Thaler. Anomalies: intertemporal choice. The journal of economic perspectives, 3(4):181–193, 1989.

45. G. J. Madden and W. K. Bickel. Impulsivity: The behavioral and neurological science of discounting. American Psychological Association, 2010.

46. J. G. March. Learning to be risk averse. Psychological review, 103(2):309, 1996.

47. J. E. Mazur. An adjusting procedure for studying delayed reinforcement. Commons, ML.; Mazur, JE.; Nevin, JA, pages 55–73, 1987.

48. S. M. McClure, K. M. Ericson, D. I. Laibson, G. Loewenstein, and J. D. Cohen. Time discounting for primary rewards. Journal of neuroscience, 27(21):5796–5804, 2007.

49. S. M. McClure, D. I. Laibson, G. Loewenstein, and J. D. Cohen. Separate neural systems value immediate and delayed monetary rewards. Science, 306(5695):503–507, 2004.

50. S. Meier and C. D. Sprenger. Temporal stability of time preferences. Review of Economics and Statistics, 97(2):273–286, 2015.

51. W. Mischel, O. Ayduk, M. G. Berman, B. Casey, I. H. Gotlib, J. Jonides, E. Kross, T. Teslovich, N. L. Wilson, V. Zayas, et al. ‘willpower’over the life span: decomposing self-regulation. Social cognitive and affective neuroscience, 6(2):252–256, 2010.

52. K. Miyazaki, K. W. Miyazaki, and K. Doya. Activation of dorsal raphe serotonin neurons underlies waiting for delayed rewards. Journal of Neuroscience, 31(2):469–479, 2011.

53. D. J. Navarick. Discounting of delayed reinforcers: measurement by questionnaires versus operant choice procedures. The Psychological Record, 54(1):85, 2004.

54. E. E. Osuna. The psychological cost of waiting. Journal of Mathematical Psychology, 29(1):82–105, 1985.

55. F. Paglieri. The costs of delay: Waiting versus postponing in intertemporal choice. Journal of the Experimental Analysis of Behavior, 99(3):362–377, 2013.

56. N. E. Paterson, C. Wetzler, A. Hackett, and T. Hanania. Impulsive action and impulsive choice are mediated by distinct neuropharmacological substrates in rat. International Journal of Neuropsychopharmacology, 15(10):1473–1487, 2012.

57. J. H. Patton, M. S. Stanford, et al. Factor structure of the barratt impulsiveness scale. Journal of clinical psychology, 51(6):768–774, 1995.

58. J. W. Peirce. Psychopy—psychophysics software in python. Journal of neuroscience methods, 162(1):8–13, 2007.

59. R. A. Poldrack, J. Clark, E. Pare-Blagoev, D. Shohamy, J. C. Moyano, C. Myers, and M. A. Gluck. Interactive memory systems in the human brain. Nature, 414(6863):546, 2001.

60. C. Prévost, M. Pessiglione, E. Météreau, M.-L. Cléry-Melin, and J.-C. Dreher. Separate valuation subsystems for delay and effort decision costs. Journal of Neuroscience, 30(42):14080–14090, 2010.

61. P. J. Reber, D. R. Gitelman, T. B. Parrish, and M. M. Mesulam. Dissociating explicit and implicit category knowledge with fmri. Journal of cognitive neuroscience, 15(4):574–583, 2003.

62. A. D. Redish, S. Jensen, and A. Johnson. Addiction as vulnerabilities in the decision process. Behavioral and Brain Sciences, 31(4):461–487, 2008.

63. E. Reuben, P. Sapienza, and L. Zingales. Time discounting for primary and monetary rewards. Economics Letters, 106(2):125–127, 2010.

64. B. Reynolds and R. Schiffbauer. Measuring state changes in human delay discounting: an experiential discounting task. Behavioural processes, 67(3):343–356, 2004.

65. A. J. Robison and E. J. Nestler. Transcriptional and epigenetic mechanisms of addiction. Nature reviews neuroscience, 12(11):623, 2011.

66. A. G. Rosati, J. R. Stevens, B. Hare, and M. D. Hauser. The evolutionary origins of human patience: temporal preferences in chimpanzees, bonobos, and human adults. Current Biology, 17(19):1663–1668, 2007.

67. P. A. Samuelson. A note on measurement of utility. The review of economic studies, 4(2):155–161, 1937.

68. P. A. Samuelson. Some implications of “linearity”. The Review of Economic Studies, 15(2):88–90, 1947.

69. S. Sanchez-Roige, P. Fontanillas, S. L. Elson, A. Pandit, E. M. Schmidt, J. R. Foerster, G. R. Abecasis, J. C. Gray, H. de Wit, L. K. Davis, et al. Genome-wide association study of delay discounting in 23,217 adult research participants of european ancestry. Nature neuroscience, 21(1):16, 2018.

70. G. Schoenbaum, M. R. Roesch, T. A. Stalnaker, and Y. K. Takahashi. A new perspective on the role of the orbitofrontal cortex in adaptive behaviour. Nature Reviews Neuroscience, 10(12):885, 2009.

71. M. S. Stanford, C. W. Mathias, D. M. Dougherty, S. L. Lake, N. E. Anderson, and J. H. Patton. Fifty years of the barratt impulsiveness scale: An update and review. Personality and individual differences, 47(5):385–395, 2009.

72. S. C. Tanaka, K. Yamada, H. Yoneda, and F. Ohtake. Neural mechanisms of gain-loss asymmetry in temporal discounting. Journal of Neuroscience, 34(16):5595–5602, 2014.

73. S. E. Tedford, A. L. Persons, and T. C. Napier. Dopaminergic lesions of the dorsolateral striatum in rats increase delay discounting in an impulsive choice task. PloS one, 10(4):e0122063, 2015.

74. R. Thaler. Some empirical evidence on dynamic inconsistency. Economics letters, 8(3):201–207, 1981.

75. A. Tymula, L. A. R. Belmaker, L. Ruderman, P. W. Glimcher, and I. Levy. Like cognitive function, decision making across the life span shows profound age-related changes. Proceedings of the National Academy of Sciences, 110(42):17143–17148, 2013.

76. A. A. Tymula and P. W. Glimcher. Expected subjective value theory (esvt): A representation of decision under risk and certainty. 2016.

77. T. W. Watts, G. J. Duncan, and H. Quan. Revisiting the marshmallow test: A conceptual replication investigating links between early delay of gratification and later outcomes. Psychological science, page 0956797618761661, 2018.

78. R. Webb, P. W. Glimcher, and K. Louie. Rationalizing context-dependent preferences: Divisive normalization and neurobiological constraints on choice. 2016.

79. A. M. Wikenheiser, D. W. Stephens, and A. D. Redish. Subjective costs drive overly patient foraging strategies in rats on an intertemporal foraging task. Proceedings of the National Academy of Sciences, 110(20):8308–8313, 2013.

80. M. Wittmann and M. P. Paulus. Temporal horizons in decision making. Journal of Neuroscience, Psychology, and Economics, 2(1):1, 2009.

81. S.-W. Wu, M. R. Delgado, and L. T. Maloney. Economic decision-making compared with an equivalent motor task. Proceedings of the National Academy of Sciences, 106(15):6088–6093, 2009.

82. H. Yamada, A. Tymula, K. Louie, and P. W. Glimcher. Thirst-dependent risk preferences in monkeys identify a primitive form of wealth. Proceedings of the National Academy of Sciences, 110(39):15788–15793, 2013.

83. G. Zauberman, B. K. Kim, S. A. Malkoc, and J. R. Bettman. Discounting time and time discounting: Subjective time perception and intertemporal preferences. Journal of Marketing Research, 46(4):543–556, 2009.

